# Ancestry-specific maps of GRCh38 linkage disequilibrium blocks for human genome research

**DOI:** 10.1101/2022.03.04.483057

**Authors:** James W. MacDonald, Tabitha A. Harrison, Theo K. Bammler, Nicholas Mancuso, Sara Lindström

## Abstract

A map of approximately independent linkage disequilibrium (LD) blocks has many uses in statistical genetics. Current publicly available LD block maps are based on sparse recombination maps and are only available for GRCh37 (hg19) and prior genome assemblies. We generated LD blocks in GRCh38 coordinates for African (AFR), East Asian (EAS), European (EUR) and South Asian (SAS) ancestry populations. These new maps consist of 1,143 (EAS) - 1,604 (AFR) independent LD blocks across the 22 autosomal chromosomes and can be accessed at https://github.com/jmacdon/LDblocks_GRCh38.

## Introduction

With an increasing number of large-scale genome-wide association studies (GWAS) relying on meta-analysis, many newly developed statistical methods circumvent the need for individual-level data, and instead require GWAS summary statistics only. These methods often rely on an external reference panel such as the 1,000 Genomes Project (1KGP) (The 1000 Genomes Project Consortium, 2015) to model patterns of population-specific linkage disequilibrium (LD), which refers to the non-random association between alleles at two loci within a population. Further, approaches to study the local genetic architecture at a specific genomic locus utilize population-specific maps of approximately independent LD blocks (Shi *et al*., 2019). Methods to build such blocks have been described previously (Berisa and Pickrell, 2016). In this context, an LD block is a genomic region with relatively strong LD between alleles, and the LD between adjacent blocks is low. Blocks are often interspersed by recombination hotspots. The ability to broadly define sets of LD blocks with the desired property of showing high within-block LD but low between-block LD is critical for many down-stream GWAS analysis tools. Current publicly available LD blocks were generated based on GRCh37 coordinates, and as more data are mapped to the GRCh38 genome assembly (including resources such as GTEx, UK Biobank and the All of Us Research Program), updated block coordinates are needed.

One potential and straightforward solution would be to convert existing GRCh37 LD blocks to GRCh38 positions using resources such as liftOver (Kent *et al*., 2002). However, this approach can produce large unmappable regions, as liftOver aims to map short genomic sequences between genome builds, with longer genomic regions becoming fragmented and scattered across multiple chromosomes. Alternatively, the GRCh37 recombination map could be lifted to GRCh38 and then used to generate LD blocks. Current GRCh37 blocks utilized a recombination map based on HapMap II data, which was originally mapped to NCBI build 35. Even though it has been shown that recombination maps lifted over from GRCh37 to GRCh38 and maps directly generated in GRCh38 are highly correlated (Spearman correlation > 0.98) (Spence and Song, 2019), repetitive lifting over of recombination maps between genome builds could introduce errors, as rearrangements between different multiple genome builds can result in SNP positions being inverted. Such SNPs would have to be manually removed, as they would introduce negative recombination rates in those regions.

In this study, we leveraged recently generated genome-wide recombination maps using population-specific GRCh38 1KGP phase 3 (Spence and Song, 2019) data to generate LD blocks for populations of African (AFR), East Asian (EAS), European (EUR), and South Asian (SAS) ancestry.

## Methods & Materials

We applied LDetect (Berisa and Pickrell, 2016) (https://bitbucket.org/nygcresearch/ldetect/src/master/), which utilizes population-specific genetic variants and a recombination map file to generate LD blocks. Briefly, the method computes the covariance matrix between variants using the 1KGP data, and then uses the Wen-Stephens shrinkage estimator (Wen and Stephens, 2010) to reduce covariance for sufficiently distant variants to zero. The method then vectorizes the covariance matrix by summing the off-diagonal covariance estimates and identifies ‘dips’ in the vectorized covariance by first using a Fourier transform to smooth the data, and then a local search algorithm to identify local minima. We used the default values recommend by the LDetect authors for each step.

We used recombination rates based on each of the 26 phase 3 1KGP populations generated by Spence and Song (Spence and Song, 2019). In the 1KGP dataset, continental superpopulation ancestry groups consist of multiple subpopulation ancestry groups. To generate LD blocks for each continental superpopulation group, we used recombination maps from each subpopulation with the largest sample size (AFR: Gambian in Western Division, the Gambia – Mandinka (n=113), GWD; EAS: Han Chinese in Beijing, China (n=106), CHB; EUR: Iberian populations in Spain (n=107), IBS; and SAS: Gujarati Indian in Houston, TX (n=113), GIH).

We downloaded VCF files from the 1KGP GRCh38 December 2018 biallelic single nucleotide variant (SNV) data (http://ftp.1000genomes.ebi.ac.uk/vol1/ftp/data_collections/1000_genomes_project/release/20181203_biallelic_SNV/). We included all 1KGP subjects within each ancestry group, with the exceptions of African Caribbean in Barbados (ACB) and African Ancestry in Southwest US (ASW) from the AFR ancestry group (due to admixture) and Finnish in Finland (FIN) from the EUR ancestry group (a genetically isolate population). The total number of subjects included per superpopulation are AFR (n=513), EAS (n=515), EUR (n=417), and SAS (n=492). We removed variants with a minor allele frequency (MAF)<0.01. We interpolated genetic distances for the remaining variants using the population-specific recombination maps and a custom R script (see GitHub repository scripts). Briefly, the recombination maps provided genomic ranges with a relatively constant recombination rate (chromosome, start, end, recombination rates (in cM/Mb), and genetic distances). We used linear interpolation of the recombination maps to estimate genetic distances for each variant in the filtered VCF files. The variant data and recombination maps were then used to compute covariance matrices between genetic variants. This is the most computationally expensive step and can be streamlined by splitting each chromosome into smaller partitions to process in parallel. Specifically, we used the default arguments for LDetect to partition each chromosome by first generating naïve, non-overlapping blocks containing 5,000 SNVs each. To ensure that these naïve blocks did not terminate within regions of high LD, each partition was extended by LDetect until a shrunken covariance estimator (Wen and Stephens, 2010) between the first and last SNV in the block was negligible (<1.5×10^−8^).

Resulting partitions and overlaps are presented in **Table 1**. LDetect computed covariance minima across each partition and selected block-specific breakpoints using a low-pass filter with local search algorithm (fourier-ls algorithm). The low-pass filter identifies LD block boundaries based on the covariance minima and an average number of SNPs per block. We set the average block size to 7,000 SNPs to obtain comparable block sizes to the previously derived GRCh37 blocks, which used 10,000 SNPs (Berisa and Pickrell, 2016), reflecting the smaller number of SNVs in our data after filtering.

**Table 1.** Partition sizes and overlaps in megabases (Mb)

|  | AFR | EAS | EUR | SAS |
| --- | --- | --- | --- | --- |
| **Median Partition Size (Mb)** | 1.69 | 1.61 | 1.60 | 1.65 |
| **Median Overlap (Mb)** | 0.56 | 0.49 | 0.47 | 0.52 |

AFR: African, EAS: Eastern Asian, EUR: European, SAS: South Asian

## Data Availability

Details, including code and generated LD block coordinates in BED format, can be found in the following GitHub repository: https://github.com/jmacdon/LDblocks_GRCh38. Additionally, a static version is available as a DOI (<u>DOI: 10.5281/zenodo.7859072</u>).

## Results and Discussion

LD block statistics by superpopulation are presented in Table 2 and Figure 1. The number of blocks genome-wide ranged between 1,143 (EAS) and 1,604 (AFR), with an average length of 1.6Mb (AFR) to 2.2Mb (EAS). We observed a higher proportion of blocks <1.5Mb in AFR compared to other populations (Figure 1), consistent with previous observations that decay in LD as a function of physical distance is fastest in African ancestry populations and slowest in East Asian populations (The 1000 Genomes Project Consortium, 2015). Compared to previous blocks, the blocks presented here are fewer and on average longer. The majority of block lengths (>97.5% in each ancestry group) were within 2 standard deviations from the population-specific mean. Blocks that contained less than 100 SNPs (two blocks for AFR), were merged into the adjacent block for usability. Adittionally, blocks that overlap any chromosome’s centromere were excluded.

**Table 2.** LD block number and length by 1KGP superpopulations

| Superpopulation | Number of blocks | Block length (Mb) |  |  |  |
| --- | --- | --- | --- | --- | --- |
|  |  | Minimum | Median | Mean | Maximum |
| AFR | 1,604 | 0.0325 | 1.58 | 1.75 | 26.5 |
| EAS | 1,143 | 0.0617 | 2.19 | 2.46 | 33.5 |
| EUR | 1,360 | 0.13 | 1.84 | 2.06 | 28.9 |
| SAS | 1,291 | 0.0229 | 1.97 | 2.17 | 29.4 |

AFR: African, EAS: Eastern Asian, EUR: European, SAS: South Asian, bp: base pair.

**Figure 1:**
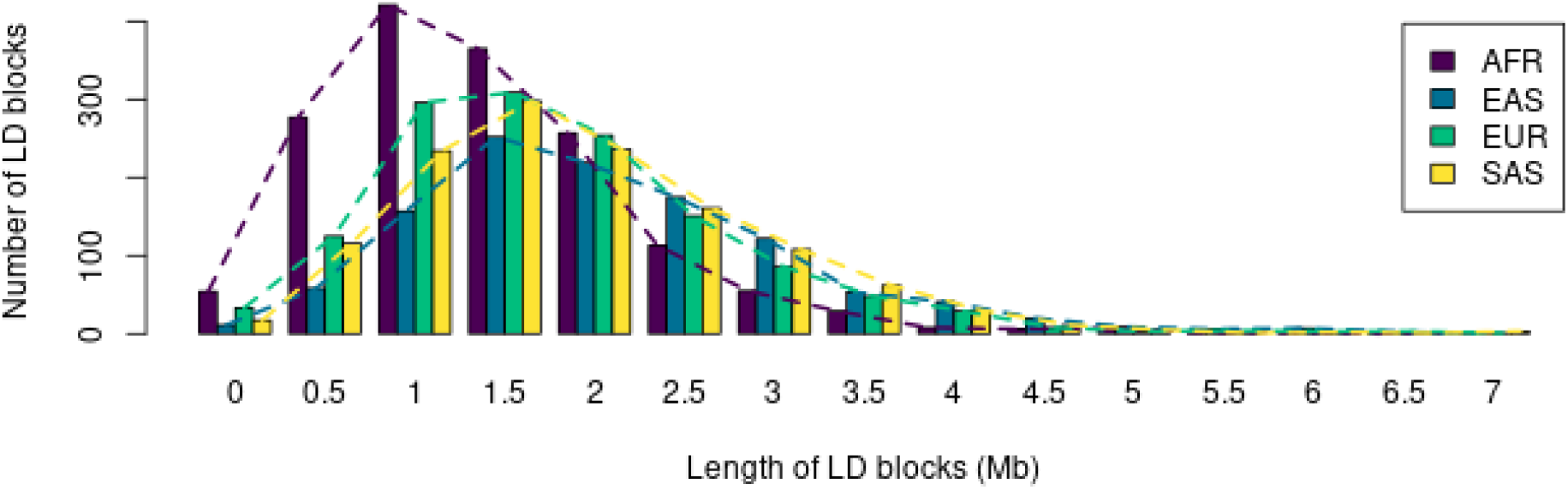
Distribution of LD block size by 1KGP superpopulation. Note that the x axis is truncated for illustrative purposes.

In this manuscript, we describe a newly developed GRCh38 map of approximately independent LD blocks for superpopulations of African, East Asian, European, and South Asian ancestry. This resource will allow researchers to leverage existing statistical methods on GRCh38 data that require information about LD block boundaries generated from the same genome build.

